# Refining gene delivery to skeletal muscle with a dual-strategy approach of muscle-tropic AAV capsids and muscle-specific promoters

**DOI:** 10.1101/2024.08.02.605568

**Authors:** Annalucia Darbey, Wenanlan Jin, Linda Greensmith, James N. Sleigh, John Counsell, Pietro Fratta

**Affiliations:** Department of Neuromuscular Diseases and UCL Queen Square Motor Neuron Disease Centre, UCL Queen Square Institute of Neurology, Queen Square, London WC1N 3BG, UK; UK Dementia Research Institute, University College London, London WC1E 6BT, UK; Research Department of Targeted Intervention, UCL Division of Surgery and Interventional Science, Charles Bell House, London, UK; The Francis Crick Institute; London, NW1 1AT, UK

**Keywords:** gene therapy, neuromuscular diseases, skeletal muscle, AAV, promoters

## Abstract

Viral vector technologies based on adeno-associated virus (AAV) have demonstrated promising ability to deliver genetic cargo to a range of organs *in vivo,* with several novel candidates showing clinical efficacy in human trials over the past decade. However, naturally occurring AAV serotypes are limited in their ability to target skeletal muscle, an important gene therapy target for many neuromuscular disorders. This means that high doses of AAV are often required to achieve therapeutically effective doses in muscle. To overcome this, novel AAV vector capsids have been engineered by inserting targeting peptides into the AAV9 capsid variable region VIII (VRIII) to achieve greater muscle transduction efficiency. Here we describe investigation of a newly reported capsid, called MyoAAV1A combined with clinically validated muscle-specific promoters. We profiled the efficiency of *in vivo* delivery to murine skeletal muscle and found that the optimal combination of MyoAAV1A capsid with MHCK7 promoter maintains transgene expression in skeletal muscle, and reduces expression in off-target tissues, particularly the liver. This highlights a promising capsid-promoter combination to progress in further preclinical research for skeletal muscle gene therapy.

**Graphical abstract:** 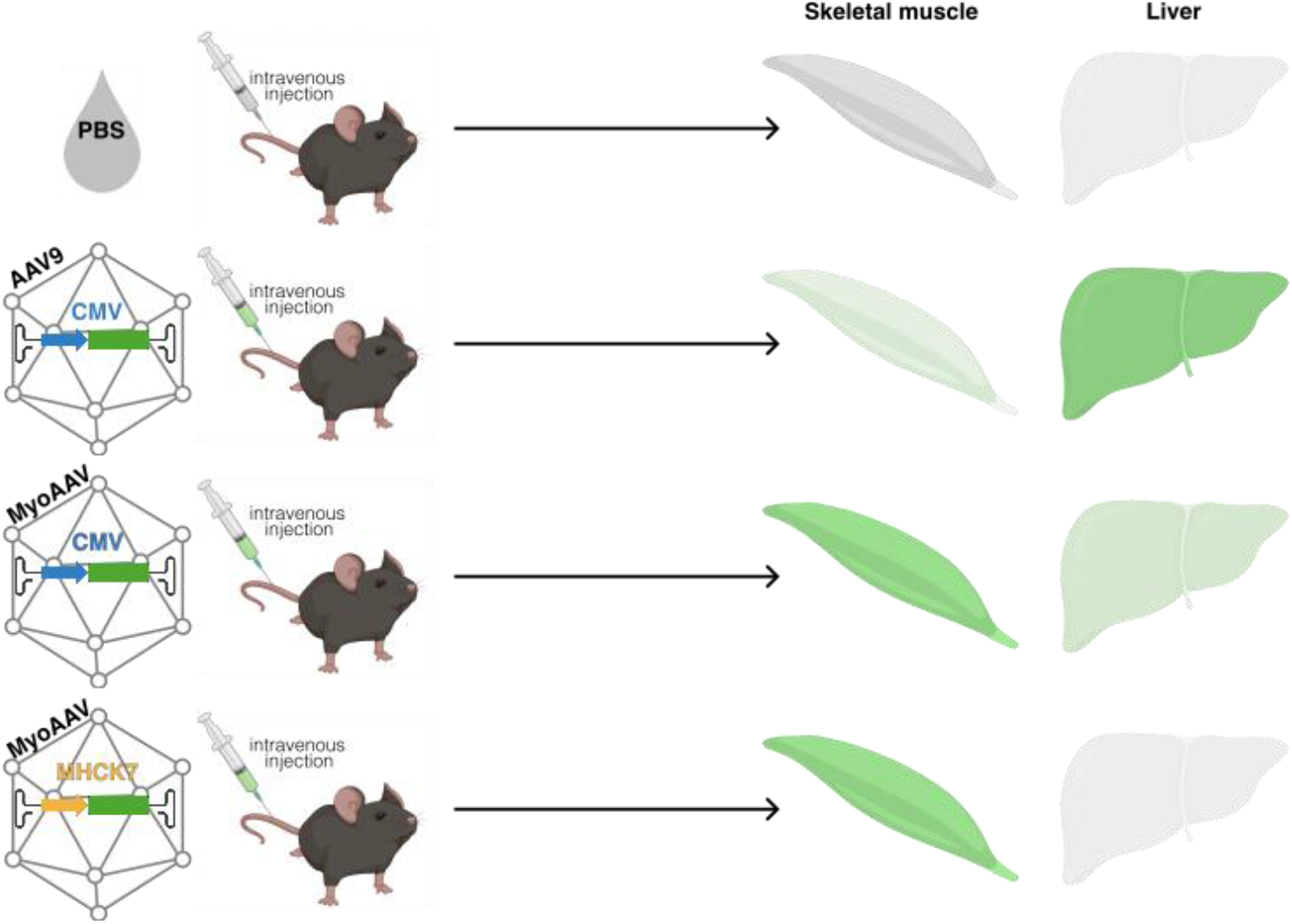

## Introduction

Recombinant adeno-associated virus (AAV) vectors^1,2^ are becoming an important delivery vehicle for gene supplementation, gene editing and gene knock-down approaches in clinical and preclinical studies, with AAVs having been or are currently being utilised in >200 clinical trials globally^3^. However, major challenges that remain with current AAV vectors are delivery efficiency, specificity and avoiding immune responses, prompting a surge in AAV capsid research aiming to modify or redirect vectors to improve selective transduction of target cells. A popular approach for AAV capsid engineering involves the insertion of peptides within the highly diverse regions of the capsid protein known as “variable regions” (VRs), which are responsible for serotype-specific preferences of receptor binding and cell transduction^4^. Due to their exposed position on the capsid surface and their function in receptor binding, the VR regions have been targets for genetic modification using strategies such as directed-evolution and rational capsid design, the former of which was utilised in the development of CNS targeting vector AAV-PHP.eB^5,6^ and muscle-tropic vectors AAVMYO^7^ and MyoAAV^8^. Delivery of high viral titres to achieve therapeutic thresholds of expression can come at a cost of increased risk of liver toxicity. Indeed, liver toxicity is a valid concern with regards to gene therapy applications requiring high doses because of deaths linked to progressive liver dysfunction in a Phase II clinical trial of high dose gene^9^. Delivery efficiency is, therefore, of particular importance when targeting tissues that require higher doses due to transduction difficulties or account for a large percentage of total body mass, such as skeletal muscle, and therefore forcibly require higher doses. Another strategy to restrict expression of delivered transgenes to target tissue is to use tissue-specific promoters to drive transgene expression. However, tissue-specific promoters are often found to be less efficacious than commonly used, ubiquitous promoters, causing concern that their use could reduce potency of delivered transgenes^10^. Here, we have demonstrated that combining an AAV engineered to preferentially transduce muscle with a muscle-specific promoter results in reduced “off-target” expression, such as in the liver, whilst maintaining the high levels of expression often required for therapeutic benefit.

## Results

### MyoAAV delivers a superior transgene payload in adult mouse skeletal muscle than AAV9

Previous studies have demonstrated preferential muscle transduction using AAVs packaged with muscle-tropic capsids identified through directed evolution^7,8^. To assess muscle tropisms of MyoAAV capsid in our system, we compared MyoAAV and AAV9 expressing GFP under the CMV promoter. We injected 8-week-old C57BL/6J mice intravenously with 1.6e+11 vg (∼6.4e+12vg/kg) AAV9-CMV-GFP or MyoAAV-CMV-GFP and analysed transgene expression and biodistribution seven days post-injection (Figure 1a). Fluorescent imaging of whole muscle showed a stronger GFP signal in skeletal muscles injected with MyoAAV-CMV-GFP compared to those injected with AAV9-CMV-GFP and this was confirmed by quantitative analysis of GFP expression by qRTPCR and western blots (Figure 1b-e). In contrast, higher levels of GFP transgene were detected in livers from AAV9-CMV-GFP-injected mice compared to those injected with MyoAAV-CMV-GFP (Figure 1b, 1c, 1f-g). Together, these observations confirm that MyoAAV preferentially transduces muscle when delivered intravenously to adult mice.

**Figure 1.**
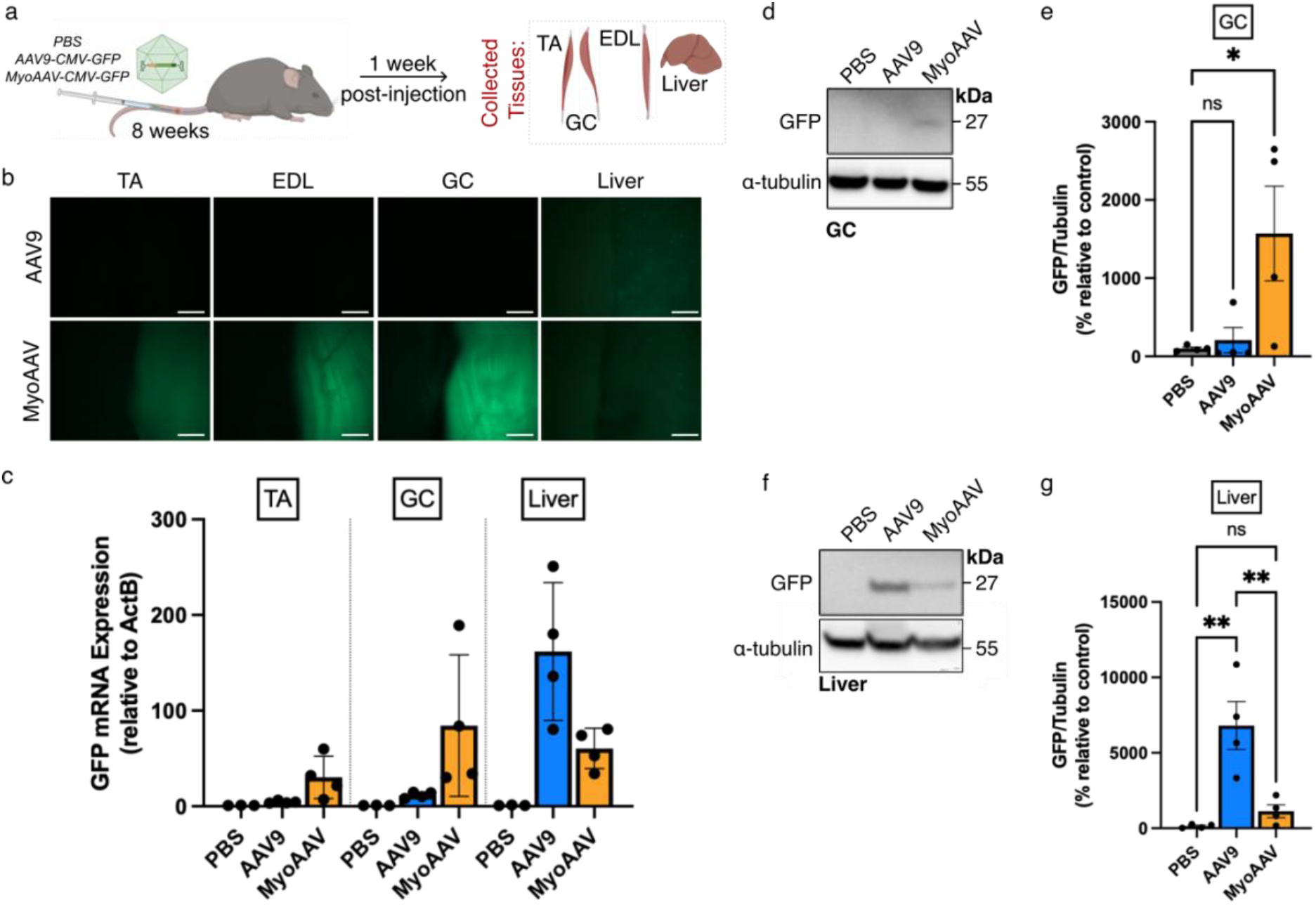
MyoAAV delivers a superior transgene payload in adult mouse skeletal muscle than AAV9. a) Schematic diagram of *in vivo* experiments performed in this study. *ITR*, inverted terminal repeat. b) Representative macroscopic images of freshly dissected tibialis anterior (TA), extensor digitorum longus (EDL), gastrocnemius (GC) muscles and liver one-week post-tail vein injection with AAV9-CMV-GFP or MyoAAV-CMV-GFP. c) *GFP* mRNA expression in TA, GC and liver relative to housekeeping gene expression (*ActB*). d&f) Western blot images and e&g) quantification of GC (d&e) and liver (f&g) one-week post-tail vein injection with AAV9-CMV-GFP or MyoAAV-CMV-GFP. α-tubulin was used as a loading control. n=3-5 animals per group. (one-way ANOVA, *P<0.05, **P<0.01, *ns* not significant). Scale bars = 200 µm.

**Figure 2.**
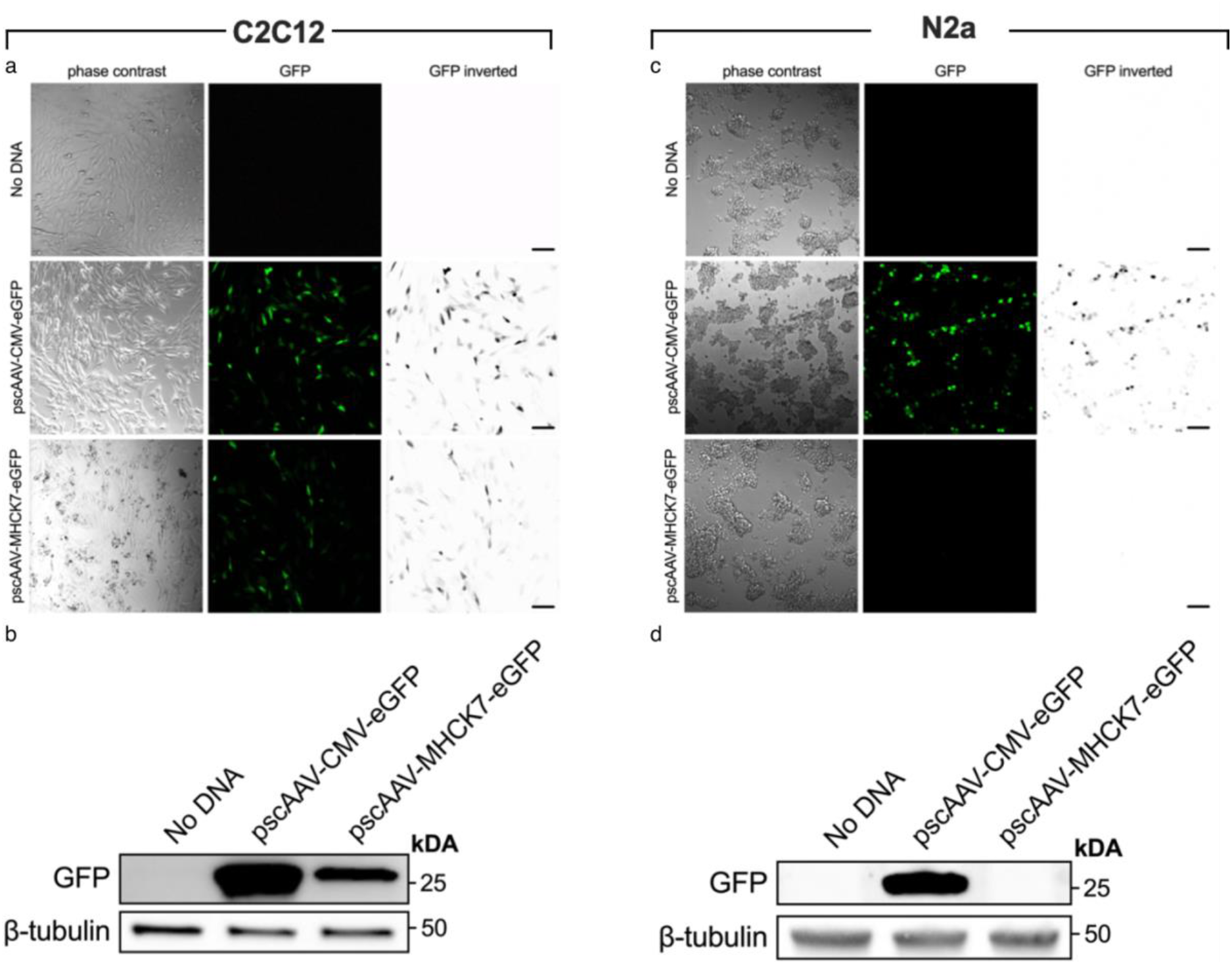
The MHCK7 promoter drives muscle cell-selective transgene expression in vitro. a) Immunofluorescence and phase contrast images of C2C12 myoblast cells magnetofected with either pscAAV-CMV-GFP or pscAAV-MHCK7-GFP. b) Western blot immunostaining demonstrating GFP expression in C2C12 cells magnetofected with pscAAV-CMV-GFP and pscAAV-MHCK7-GFP. c) Immunofluorescence and phase contrast images of N2a neuroblastoma cells magnetofected with either pscAAV-CMV-GFP or pscAAV-MHCK7-GFP. d) Western blot immunostaining demonstrating GFP expression in N2a cells magnetofected with pscAAV-CMV-GFP, but not pscAAV-MHCK7-GFP. Scale bars = 100 µm.

### The MHCK7 promoter drives muscle cell-selective transgene expression in vitro

Despite reduced liver targeting by MyoAAV, expression of the GFP transgene mRNA and protein were still detectable in livers from MyoAAV-CMV-GFP-injected mice. Given the risks of toxicity associated with AAV transduction in the liver, we sought to further increase muscle-specificity of transgene expression by incorporating a muscle-specific promoter (MHCK7^11^) into our AAV plasmid. We first assessed specificity using an *in vitro* system. pscAAV-CMV-GFP and ps-AAV-MHCK7-GFP were magnetofected into myoblast cell line C2C12 and neuroblast cell line N2a. CMV promoter led to production of GFP in both cell types, but GFP expression under MHCK7 was observable in C2C12 cells but not N2a cells (Figure c&d), confirming muscle cell specificity *in vitro*.

### MHCK7 Maintains Transgene Expression in Skeletal Muscle & Reduces Expression in Liver

CMV-GFP or MHCK7-GFP plasmids were then packaged into MyoAAV capsids to generate a combined muscle-targeting approach. Vectors were injected intravenously into 8-week-old male C57BL/6J mice at high (1e+12vg total) and low (1.92e+11vg total) doses. Tissue-or cell-specific promoters have previously been reported to result in weakened expression of downstream transgenes compared to ubiquitously expressing promoters due to reduced transcriptional activity^10^. However, our fluorescent imaging and mRNA analysis of skeletal muscle revealed comparable levels of GFP transgene expression in skeletal muscles of mice injected with MyoAAV-CMV-GFP and MyoAAV-MHCK7-GFP (Figure 3a). When driven by a muscle-specific promoter, transgene mRNA and protein expression in liver were reduced to a significantly lower level compared to transgenes delivered by the ubiquitous CMV promoter. This level was not significantly different from PBS-injected control (Figure 3c, f & j). In conclusion, the MHCK7 promoter in combination with a MyoAAV capsid enhances detargeting of liver, further improving the safety profile of this muscle-targeting strategy.

**Figure 3.**
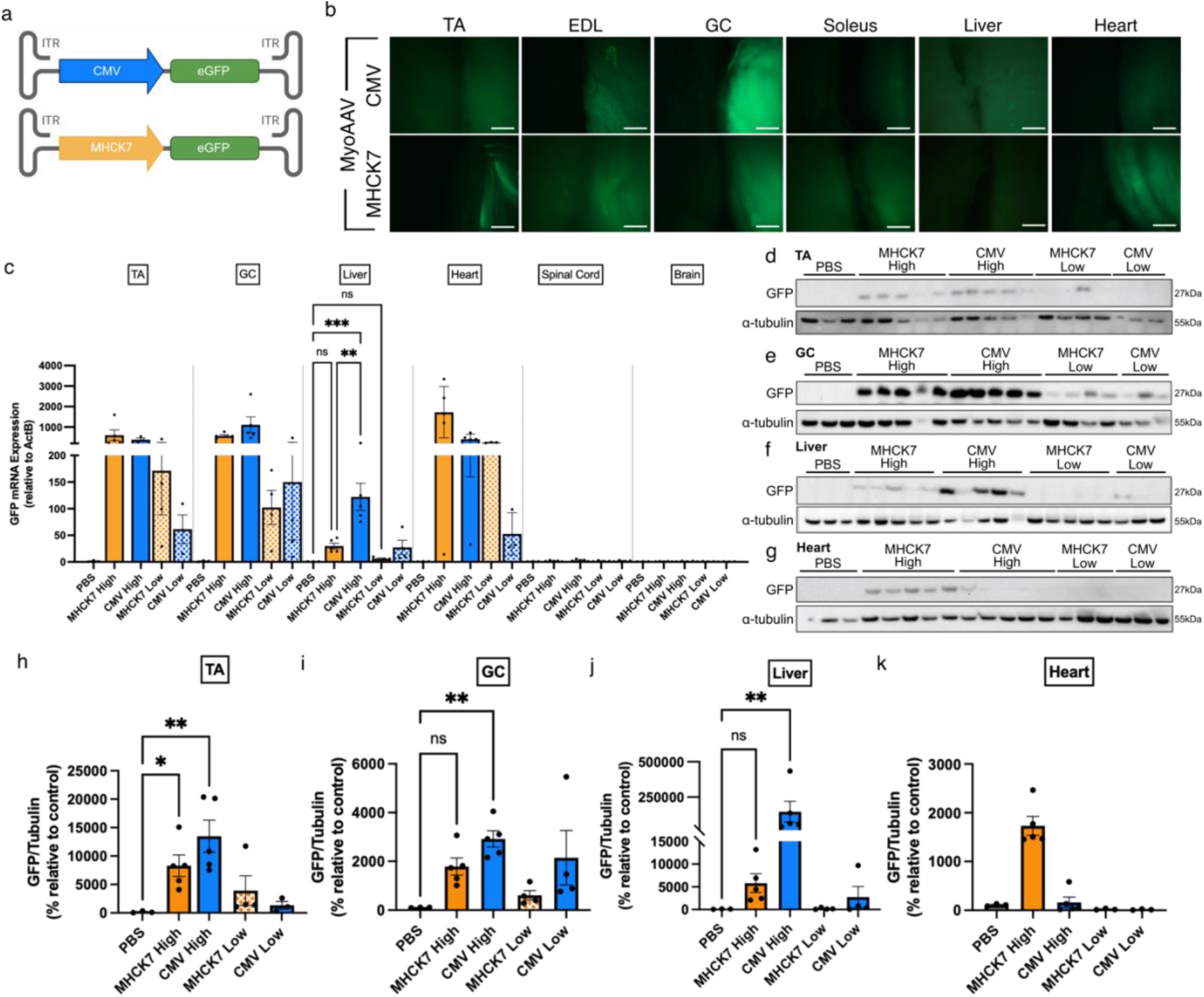
MHCK7 Maintains Transgene Expression in Skeletal Muscle & Reduces Expression in Liver. a) Schematic diagrams of AAV transgenes delivered: MyoAAV-CMV-GFP and MyoAAV-MHCK7-GFP. b) Representative macroscopic images of freshly dissected tibialis anterior (TA), extensor digitorum longus (EDL), gastrocnemius (GC), soleus muscles, liver and heart one-week post-tail vein injection with either MyoAAV-CMV-GFP or MyoAAV-MHCK7-GFP. Images taken are from the “high dose” groups injected with a total of 1e+12 vg c) *GFP* mRNA expression in collected TA, GC, liver, heart, spinal cord and brain one-week post-injection with either MyoAAV-CMV-GFP or MyoAAV-MHCK7-GFP at high (1e+12 vg) and low (1.92e+11 vg) doses. mRNA expression is relative to housekeeping gene expression (*ActB*). Western immunoblots of d) TA, e) GC, f) liver and g) heart one-week post-injection with either MyoAAV-CMV-GFP or MyoAAV-MHCK7-GFP at high and low doses. Quantification of western blots of h) TA, i) GC, j) liver and k) heart relative to α-tubulin loading controls. n=3-5 animals per group. (one-way ANOVA, *P<0.05,**P<0.01, ***P<0.001, *ns* not significant). Scale bars = 200 µm.

## Discussion

AAVs are increasingly becoming the vector of choice for *in vivo* gene therapy approaches. However, despite the AAV safety features of low integration and immunogenicity, risk of dose-associated liver toxicity remains a concern. To circumnavigate this, focus has shifted from the use of naturally occurring AAV serotypes, with a wide range of tissue tropisms, to AAV vectors with capsids engineered for cell-/tissue-specific preferences. This reduces the delivered titres required to achieve therapeutically relevant levels of expression in target tissues, preventing sequestration of viral vectors in the liver and thus lowering the likelihood of liver toxicity. This is of particular importance when the target tissue(s) cover a large expanse of the body. For example, skeletal muscle is a primary tissue affected in several neuromuscular diseases such as Duchenne Muscular Dystrophy, that requires high doses for comprehensive targeting. Recent progress in the development of vectors with preferential tissue transduction has resulted in identification of two RGD motif-containing muscle-tropic capsids, known as AAVMYO and MyoAAV^7,8^. This RGD motif is a known binding partner for integrins, suggesting that these capsids are interacting with and binding to specific integrin heterodimers on the surface of cells preferentially transduced by these vectors such as those found in skeletal muscle.

In this work, we first confirmed expression of transgenes in skeletal muscle when delivered systemically via intravenous injection of MyoAAV. We found significantly increased expression of transgenes in skeletal muscles of animals injected with MyoAAV compared to those injected with AAV9. Though shown to limit sequestration of vectors in the liver when compared to naturally occurring serotype AAV9^8^, we found transgene expression levels in liver to be comparable to those in muscle following intravenous delivery at higher doses. Therefore, despite significant muscle tropism, use of capsids with preferences to transduce skeletal muscle alone may not be sufficient to avoid liver toxicity. We therefore sought to identify a strategy in which we could maintain transgene expression in target tissues whilst reducing expression in liver. This is of particular importance given previous reports of deaths as a result of liver damage following high-dose gene therapy in a phase 2 clinical trial^9^. By utilising a muscle-specific promoter to drive transgene expression in combination with a muscle-tropic capsid, we have demonstrated a significant reduction of transgene expression in the liver whilst maintaining high levels of skeletal muscle expression, even when delivered at higher doses.

Given the limited availability of promoters that are both efficient and specific to target-tissues, it may also be paramount to incorporate other strategies of tissue-targeting or de-targeting to achieve a strong/therapeutically relevant level of transgene expression only in tissues of interest. Other methods of modulating viral transgene expression in a cell specific manner, include the use of endogenous micro RNAs (miRNAs) to silence delivered transgene expression in non-target tissues^12^ and altering endogenous expression of receptors required for viral vector uptake^13–15^.

Drawing from our data, it is evident that a combination of approaches, such as our dual strategy of using muscle-tropic capsids with muscle-specific promoters, offers a promising way forward to achieve therapeutically relevant levels of transgene expression in target tissue whilst limiting transgene expression in non-target tissues, particularly the liver. Thus, by utilising multiple strategies to tightly control expression of delivered gene therapies, it may be possible to significantly limit off-target effects, increasing the safety and effectiveness of gene therapy in clinical applications.

## Materials & Methods

### Mice

All experimental procedures were carried out under licence from the UK Home Office (Scientific Procedures Act 1986) and following approval by the UCL Institute of Neurology Animal Welfare Ethical Review Panel. 7-week-old male C57Bl6/J mice were purchased from Envigo and allowed to acclimatise for 1 week prior to any procedures taking place. Experiments were designed to use the minimum number of animals required for reliable statistical analysis and the 3R principles were implemented at each stage of the experimental planning. Sample sizes were chosen based on power calculations, preliminary data and previously reported data.

### pscAAV Plasmid Production

Self-complementary AAV (scAAV) expression plasmids pscAAV-CMV-eGFP and pscAAV-MHCK7-eGFP were created by OXGENE. Standard 571 bp CMV promoter and 714 bp eGFP sequences were used. The 770 bp MHCK7 promoter sequence was identified in Madden *et al* ^16^ and https://www.addgene.org/65042.

### Vector Production

AAV vectors were produced by triple transduction of HEK-293T cells essentially as described in Challis et al.^17^, with the following exceptions: cells were grown in 2-4×15 cm dishes and virus was centrifuged after the PEG precipitation and resuspended in up to 4mL chloroform. Samples were then vortexed for 2mins and centrifuged at 3000 × g for 20 min. The aqueous layer was then loaded on iodixanol gradients in a Type 70.1 rotor and centrifuged for 2 h at 52,000RPM. The AAV sample was collected, and buffer exchanged with 5% sorbitol and 0.25M NaCl in 1× PBS.

### In Vivo Delivery of Vectors, Tissue Collection & Imaging

Viral vectors were diluted in sterile PBS and delivered intravenously via a 100µL tail vein injection. AAVs were injected at either low dose (1.92e+11vg) or high dose (1e+12vg). Tissues were collected one-week post-injection and imaged in cold PBS using a Leitz DM RB microscope (Leica) under a brightfield and GFP lamp. Tissues were then snap-frozen and stored at -80°C until processing for downstream analysis of mRNA and proteins.

### Analysis of mRNA Expression

Total RNA was extracted from all mouse tissues using Qiazol lysis reagent (Qiagen, 79306) and cleaned up using a RNeasy Mini kit (Qiagen, 74104). cDNA synthesis was performed using the RevertAid First Strand cDNA Synthesis Kit (Thermofisher, K1621). A Taqman assay was used to analyse eGFP expression in tissues. The sequences of primers and probes used to analyse eGFP expression were as follows: Forward: 5’-CCACATGAAGCAGCACGACTT-3’, Reverse: 5’-GGTGCGCTCCTGGACGTA-3’ and Probe: 5’-Fam-TTCAAGTCCGCCATGCCCGAA-Tamra-3’. Pre-designed taqman assay for mouse β-actin (*ActB)* mRNA was utilised as the housekeeping control (Assay ID: Mm00607939_s1). Analyses were performed in triplicate using 5ng aliquots of cDNA on a QuantStudio 5 PCR system (Applied Biosystems). Relative expression levels were calculated by comparison with mouse β-actin expression.

### Western Blotting

Frozen tissues were homogenised in RIPA buffer containing complete protease inhibitor cocktail (Roche) using a motor homogenizer (TH115, OMNI). Lysates were incubated on a rotator at 4°C for 1h and then pre-cleared by centrifugation at 13,000 × g for 10-20mins at 4°C. Protein concentration was determined by bicinchoninic acid protein assay (Pierce). The following antibodies were used for detection in mouse tissues: anti-GFP (Sigma-Aldrich, SAB4301138, 1/1,000, RRID:AB_2750576) and anti-α-tubulin (Sigma-Aldrich, T9026, 1/1,000, RRID:RRID:AB_477593). Cells were collected and lysed in NP40 lysis buffer with freshly added protease/phosphatase inhibitor cocktail (78445, Thermo Scientific). The following primary antibodies were used: anti-GFP (Santa Cruz, sc-9996, 1/5,000, RRID:AB_627695) and anti-β-tubulin (Abcam, ab15568, 1/5,000, RRID:AB_2210952).

### Cell Culture

C2C12 (CRL-1772, ATCC) and Neuro-2a (N2a, CCL-131, ATCC) cells were maintained in Dulbecco’s modified Eagle’s medium (DMEM) with high glucose and pyruvate (11995065, Thermo Fisher) supplemented with 10% (v/v) heat-inactivated fetal bovine serum (10270, Thermo Fisher) and 1% (v/v) penicillin-streptomycin (15140122, Thermo Fisher). Cells were maintained in T75 flasks (156800, Thermo Fisher) at 37°C in a 5% (v/v) CO2 humidified atmosphere and passaged every 3-5 days when ≈70-90% confluent.

### Magnetofection & Cell Immunofluorescence

Cells were plated at an optimised density of 5e+4 cells per well in 24-well plates in 1 ml of DMEM. After 18 ±1 h, cells were magnetofected with plasmid DNA, similar to previously described^18^. For each well, an optimised ratio of 1 μg plasmid DNA to 3 µl NeuroMag (NM50200, Oz Biosciences) was combined in 100 µl Opti-MEM (11058021, Thermo Fisher) for 20 min at 37°C. 600 µl DMEM was removed from each well of cells prior to applying 100 µl plasmid-NeuroMag mix. Once all wells were treated in this manner, the plate was gently mixed and left on a magnetic plate (Oz Biosciences) for 35 min at 37°C. The magnet was then removed and the cells kept at 37°C for 24 ±1 h. Live cells were imaged using an inverted LSM780 laser-scanning microscope (ZEISS) within an environmental chamber prewarmed to 37°C.

### Statistical Analysis

Using Prism v9.5.0 (Graphpad Software, LLC), one-way ANOVAs or Kruskal-Wallis tests (non-parametric data) with subsequent post-hoc tests were performed to compare datasets. A P-value of p < 0.05 was considered statistically significant. Means ± standard error of the mean are plotted for all graphs.

## Data Availability Statement

AAV expression plasmids are covered by an MTA with OXGENE (UK). Material requests should be addressed to the corresponding author.

## Acknowledgements

We thank Molly Strom and Ana Cunha of the Crick Institute viral vector core for producing the AAV viral vectors.

## Author Contributions

AD, LG, JNS, JC and PF designed the research; AD, WJ and JNS performed experiments and analysed data. AD, JC and PF drafted the manuscript. All authors reviewed the data and revised the manuscript before submission.

